# Microbial Feast or Famine: dietary carbohydrate composition and gut microbiota metabolic function

**DOI:** 10.1101/2025.10.27.684932

**Authors:** Blake Dirks, Alex E. Mohr, Karen D. Corbin, Elvis A. Carnero, Steven R. Smith, Corrie M. Whisner, Bruce E. Rittmann, Rosa Krajmalnik-Brown

## Abstract

Diet composition plays a major role in shaping the structure and function of the gut microbiota and influencing host health. While numerous studies have investigated the impact of macronutrient type and quantity on microbiota using in vitro systems, animal models, and human participants, most of these studies focused primarily on microbial-community composition and lacked the functional information that can be gained from transcript-level analyses. In this exploratory analysis, we use metatranscriptomic data to gain a functional perspective on how dietary composition is associated with the gut microbiota and hypothesized implications for host physiology. Data were derived from a tightly controlled, randomized cross-over feeding study conducted in a metabolic ward, where participants consumed two isocaloric and eucaloric diets differing in food processing and fiber content: A Western Diet (WD) limited in fiber, resistant starch, and whole foods and a Microbiome Enhancer Diet (MBD) composed of fiber-rich, whole foods. Our prior findings showed that a WD lead to a resource-limited microbiota enriched in mucin-degrading bacteria that resorted to metabolizing host-derived organic material, while the MBD supported a resource-replete microbiota that primarily metabolized dietary fiber. The objective of this work was to explore these findings more deeply using bioinformatic analyses of metatranscriptomic data. Our analysis showed increased transcription of fiber-degrading enzymes in the MBD and mucin-degrading enzymes in the WD. While in this analysis functional diversity of the gut microbiome was not affected, differences in resistant-starch and fiber content shifted the types of metabolic processes being actively transcribed. The MBD promoted biosynthetic and carbohydrate-fermenting pathways, while the WD was characterized by enzymes for host-glycan and protein degradation. Furthermore, the MBD-supported ecosystem benefits host health via enhanced SCFA production and reduced reliance on host glycan degradation. The WD fostered increased mucin and protein breakdown pathways that yield metabolites that may harm the gut barrier and systemic metabolism.

## Introduction

The gut microbiota, a complex community of microorganisms residing in the human gastrointestinal (GI) tract, is increasingly recognized as a critical modulator of host health [1]. This microbial community plays a pivotal role in numerous physiological processes, including digestion, immune function, and the synthesis of essential nutrients and bioactive compounds [1]. The gut microbiota’s connection to a broad spectrum of health outcomes, including obesity [2], diabetes [3], and inflammatory diseases, [4] is well-documented. Importantly, environmental exposures have emerged as principal influencers to the gut microbiota [5], particularly diet, which can condition the composition and functionality of the microbiota in the short- and long-term [6–8].

Diet-associated functions playing an important role in host health include genes encoding glycosaminoglycan degradation, the production of short-chain fatty acids (SCFAs) via fermentation of complex polysaccharides, synthesis of specific lipopolysaccharides (LPS), and the biosynthesis of some essential amino acids and vitamins [9,10]. Dietary components provide exogenous substrates that fuel microbial metabolism and shape the structure of the gut microbial community in ways that are associated with host health. The mammalian gut, by sensing nutrients and microbial fermentation products, is part of the larger enteroendocrine system that plays a key role in maintenance of energy homeostasis [11]. Thus, recognizing which dietary patterns promote a health-enhancing microbiome has the potential to be used to help define practical dietary recommendations for health promotion and disease prevention [12].

Limited consumption of high-fiber, whole foods significantly affects the amount and type of substrates reaching the lower regions of the GI tract, where most microbial biomass resides [13– 15]. Western diets that are typically consumed in the US lack fiber and tend to include more processed foods [16,17]. The average adult man and woman in the US only consume around 18 g and 15 g of fiber per day, respectively, far below the are recommended 34 g and 25 g of fiber per day [18]. Thus, the actual diets are composed primarily of dietary substrates that are absorbed in the upper GI tract, potentially “starving” the colonic microbiota and promoting endogenous substrate consumption, such as host-derived mucin [19]. This is of great public-health concern, as fiber recommendations are rarely met in the US [16,17]. In addition, processed foods are estimated to comprise more than 50% of the average American diet [20], and world-wide consumption of processed foods continues to increase [21]. In contrast, diets rich in high fiber, whole foods deliver carbon and energy substrates (e.g., microbiota-accessible carbohydrates [MACs]) to the colon, which prevent gut-mucus depletion [22], encroachment of bacteria into the mucus layer [23], and downstream inflammation [24]. A high-fiber diet fosters a more diverse and metabolically active microbiota [25]. These factors underscore a critical link between diet composition and microbial health, since substrate availability in the colon profoundly influences microbial activity and, consequently, host health.

Prior controlled-feeding studies have demonstrated that high-fiber diets are associated with reduced host metabolizable energy [26,27] and that varying dietary composition can alter energy harvest efficiency in a way that correlates to community shifts in the gut microbiota [27,28]. Diets high in complex polysaccharides have resulted in altered gut microbiota linked with increased fecal, serum, or urine concentrations of SCFAs, weight loss, and improvements of cytokine and metabolome profiles [29–32]. Given that humans produce limited carbohydrate-degradation enzymes (CAZymes), the gut microbiota are required to metabolize several dietary fibers [25]. Indeed, a diet low in fiber is associated with a reduced CAZyme reservoir within the gut microbiota [33].

Although carbohydrates are the preferred substrate for microbiota, proteins are also an important source for nutrients. Protein is generally degraded into amino acids that are then fermented in the distal colon after accessible carbohydrates have been consumed [34]. Amino-acid fermentation produces metabolites such as ammonium, hydrogen sulfide, indoles, phenols, and amines that are detrimental for gut health [35]. Many experiments have demonstrated these detrimental health effects using CaCo-2 enterocyte cell lines. For example, ammonium inhibited oxidation of SCFAs in mitochondria, reduced absorptive capacity, and disrupted cell-cell adhesion [36]. Indoles affected cellular respiration, increased production of inflammatory cytokines, and damaged DNA [37]. Certain phenolic compounds, such as p-cresol, also disrupted cell adhesion and increased gut permeability [38]. Hydrogen sulfide, strongly associated with colorectal cancer, impaired butyrate oxidation, and contributed to inflammation, and is strongly associated with colorectal cancer [39]. Higher concentrations of these uremic toxins were also observed in humans with very low bowel-movement frequency which is linked to many bowel diseases [40]. High-protein, low-carbohydrate diets also alter gut microbiota in mice by increasing the abundance of pathogenic bacteria, such as *Shigella, Enterococcus*, and *Streptococcus*, while reducing the abundance of healthy bacteria, such as *Ruminococcus, Akkermansia*, and *Faecalibacteria* [41].

Diet-induced changes in gut microbial composition are well established, but the functional activities of these communities are less frequently measured directly; most studies infer metabolism from taxonomic or gene-content data [42]. Metatranscriptomics complements metagenomics by quantifying active gene expression rather than potential, enabling a dynamic approximate view of microbial responses to diet and environment [43]. Leveraging our previous work that provided evidence for interactions among diet, host, and microbial composition on energy balance [27], here we delve deeper into the changes in microbial functional activity under diets with specific macronutrient composition.

In this exploratory work, we provide metatranscriptomic evidence that the Western Diet (WD), designed to be easily absorbable by the host, and the Microbiome Enhancer Diet (MBD), designed to be minimally absorbable by the host, lead to divergent microbial-community functions. The microbiota under the WD reflected a community more reliant on host-derived substrates, consistent with a resource-limited state (“starving”), whereas under the MBD the microbiota reflected a community with a dietary fiber–driven metabolism and broader anabolic activity (“thriving”).

## Methods

### Clinical Trial

The design details of the parent clinical study (NCT02939703) from which the samples and data for this manuscript were derived are published [27,44]. Briefly, the study was approved by the AdventHealth Institutional Review Board and conducted at AdventHealth Translational Research Institute in Orlando, Florida. After signing informed consent and meeting eligibility criteria, 17 participants, 9 men and 8 women, were included in the study. The study was a randomized crossover study with a Western Diet (WD) as a control and a Microbiome Enhancer Diet (MBD) as an intervention. This design minimized the impact of confounders and the interindividual variability in the gut microbiota because each participant served as their own control. Participants were randomly assigned a diet order. Eight participants started with the WD followed by the MBD, and nine participants started with the MBD followed by the WD. Meals were uniquely prepared for each participant’s caloric needs to maintain energy balance (eucaloric). The two diets were also equivalent in energy (isocaloric) and the proportions of carbohydrates, protein and fat were the same. The diets were designed to differ by 4 components: fiber, resistant starch, processing, and food particle size [27,44]. Diets were consumed outpatient for 11 days and while domiciled in a metabolic ward for 11 days with a 14-day washout between diets. All samples and data were collected during the domiciled period. During each diet period, participant energy expenditure was measured in a whole room calorimeter for 6 days. During this time participant physical activity was tightly controlled, and fecal samples were collected.

### RNA Sequencing and sequence processing

We used RNA sequences that were previously published [45]. RNA-sequences were quality controlled using FastQC (version 0.12.0) [46]. Adapters were trimmed using TrimGalore (version 0.6.5) [47]. RNA sequences were aligned against Hg38 (GRCh38.p14) using STAR (version 2.7.11a) [48]. Aligned sequences were removed, and the remaining reads were paired and annotated with Enzyme Commission (EC) numbers using HUMAnN3 (version 3.8) [49] with standard parameters.

### Transcript alpha- and beta-diversity

All calculations and analyses were conducted using R (Version 4.2.2) [50]. EC-annotated metatranscriptomic output from HUMAnN3 (version 3.8) was processed for alpha- and beta-diversity analysis using the “phyloseq” R package (Version 1.50.0) [51]. Alpha- and beta-diversity were calculated using raw count data. Alpha-diversity metrics, including feature count, evenness, Shannon diversity, and Inverse-Simpson diversity, were calculated using the “microbiome” R package (Version 1.28.0) [52]. Bray–Curtis and Jaccard distance matrices were calculated using vegan (Version 2.7-1) [53]. The distance matrices were tested for significance by PERMANOVA using vegan (Version 2.7-1). Beta-dispersion was calculated, and the results tested for significance with PERMANOVA and Tukey’s HSD in vegan (Version 2.7-1). Figures for alpha- and beta-diversity were created using the “ggplot2” R package (Version 4.0.0).

### Differential Gene and Transcript Abundance Analysis

Genomic differential abundance and expression testing by diet were carried out using the Enzyme Commission (EC) annotated metatranscriptomic output from HUMAnN3 (version 3.8) using the R package MaAslin2 (version 1.7.3) [54]. A small pseudo-count (equal to half of the lowest non-zero count) was added to any of genes and transcripts with an abundance of zero.

The counts were then transformed by center log ratio as recommended for analysis of compositional data [55,56]. Transcripts with less than 10% prevalence and low variance (less than half of the median variance [54,57]) were then filtered to reduce the number of necessary comparisons.. A linear mixed model (LMM) was then used to determine differential transcript abundances. The model included diet, sequence, and period as fixed factors with participant as the random factor [27]. Carryover between diets is unlikely due to the tight control of the diet and environmental conditions in the whole-room calorimeter. A P value < 0.05 was considered statistically significant. When P values required correction for multiple comparisons, the Benjamini-Hochberg (BH) method was used. An adjusted P value (adj.P) < 0.25 was the exploratory threshold for noteworthy features.

### Metabolic Pathway Reconstruction

We reconstructed microbiota MetaCyc metabolic pathways in each diet using differentially abundant transcripts with the program MinPath (version 1.6) [58]. Briefly, once differentially abundant transcripts were identified in each diet, the EC number for those transcripts were then input into MinPath, which then found the minimum number of MetaCyc pathways that explain what the input transcripts are.

## Results and Discussion

### Microbiota Transcripts α- and β-diversity were not significantly different between diets

In our prior metagenomic analysis, significant diet-induced changes occurred in the structures of the microbial communities. In particular, butyrate-producers and fiber degraders increased in the MBD, while acetate-producers and simple-sugar degraders increased in the WD [27]. To determine whether these previously identified community changes were reflected in the microbial functional profile, we investigated the α- and β-diversity of the annotated transcripts, with the results summarized in **Figure 1**. In total, HUMAnN3 annotated 2,448 microbial gene transcripts that encode enzymes across both diets, with 2250 transcripts encoding enzymes in the WD and 2318 transcripts encoding enzymes in the MBD (**Figure 1A**). Both diets had 2190 transcripts encoding enzymes (89.5%) in common, while 130 (5.3%) and 128 (5.2%) transcripts encoding enzymes were unique to the WD and MBD, respectively (**Figure 1B**). Despite varying dietary inputs, α-diversity metrics, including observed features, evenness, Shannon index, and Inverse Simpson index (**Supplementary Figure 1**), did not show significant differences in the diversity of transcripts between the WD and MBD. Similarly, β-diversity analyses using Bray-Curtis dissimilarity and Jaccard index did not show significant separation based on (P ≥ 0.136, P ≥ 0.169; **Figure 1C**).

**Figure 1.**
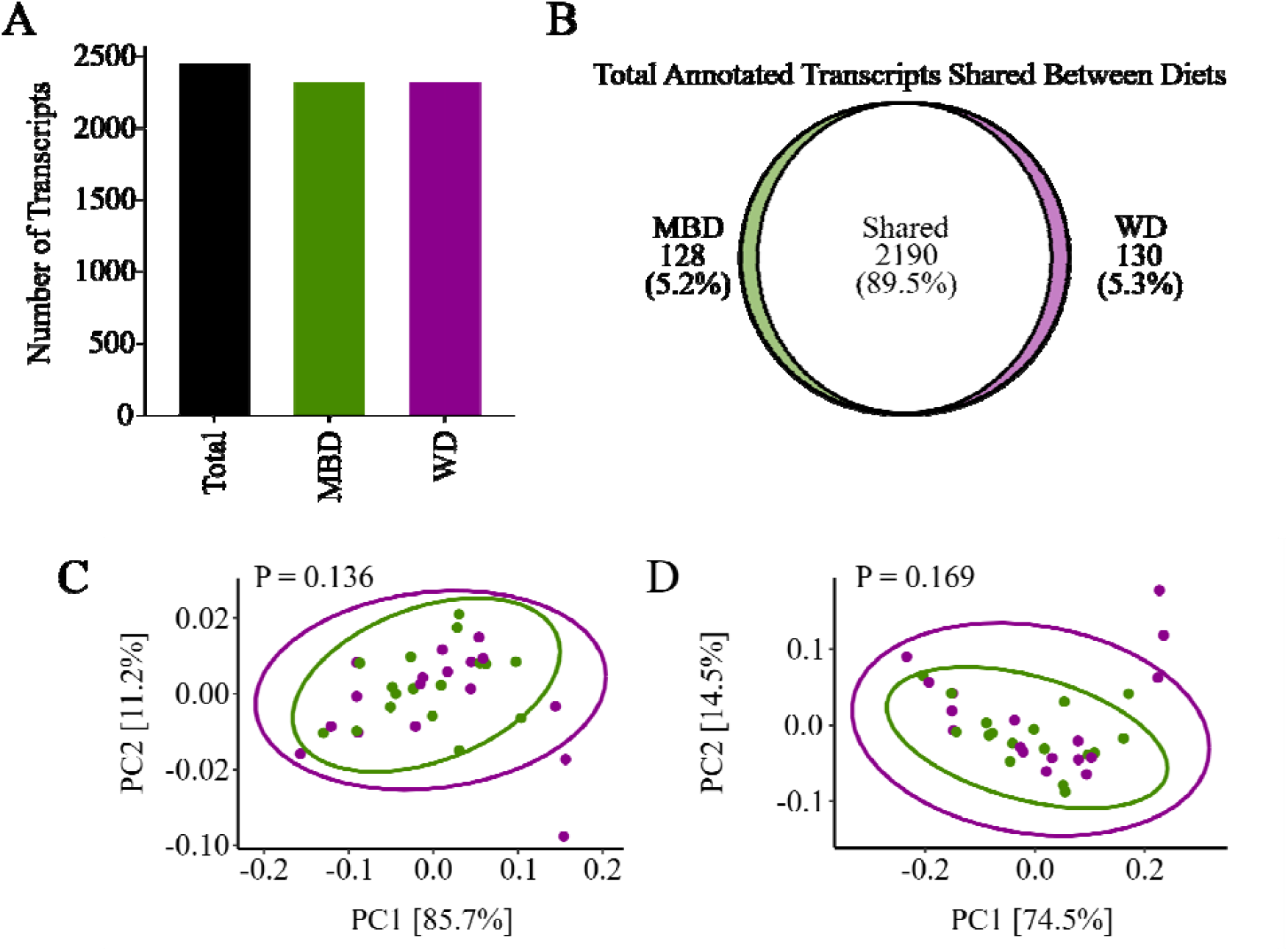
Total transcripts detected and β-diversity were not significantly different between diets. A) Total detected features (2,448) and detected features in the MBD (2,318) and WD (2,250). B) Shared and unique features between the MBD and WD. C) Bray-Curtis (Dis)similarity for transcripts was not significantly different between diets. D) Jaccard Index for transcripts was not significantly different between diets. Bray-Curtis and Jaccard metrics were tested by analysis of similarity (PERMANOVA). N = 17, MBD: Microbiome Enhancer Diet, WD: Western Diet.

The similarity of α- and β-diversity metrics of transcripts encoding enzymes between diets may be due to functional redundancy, a core characteristic of the gut microbiota [59]. This functional redundancy usually reflects the gut community’s resilience to shifts in substrate availably by maintaining broad metabolic capabilities because it can modify gene expression among existing community members [60]. Such ecological stability in microbial transcriptomes also has been documented in other dietary-intervention and observational studies, where overall functional diversity remained relatively steady despite considerable shifts in nutrient composition [61,62]. While this finding might suggest limited impacts of diet on overall community stability [63–65], our subsequent analyses clearly demonstrate that diet-induced shifts occurred robustly at the level of specific functional activities, but not through aggregated functional diversity.

### Microbiota in the MBD shifted its metabolism towards growth and degradation of diet-derived nutrients, while in the WD shifted towards amino acid degradation

To explore further differences in microbial functional activity between diets, we reconstructed biosynthetic and degradative MetaCyc pathways from differentially abundant transcripts. First, we filtered transcripts for low prevalence and variance and then performed differential-abundance analysis on the remaining 2053 transcripts using MaAsLin2 [54]. After we applied the Benjamini-Hochberg method to correct for multiple comparisons (FDR ≤ 0.25 as the exploratory threshold), 454 transcripts were differentially abundant by diet, with 331 transcripts more abundant on the MBD and 123 transcripts more abundant on the WD (**Figure 2A**). Using these differentially abundant transcripts, we reconstructed MetaCyc metabolic pathways in each diet using MinPath [58], a tool that finds the minimum number of MetaCyc pathways that contain all the input transcripts.

**Figure 2.**
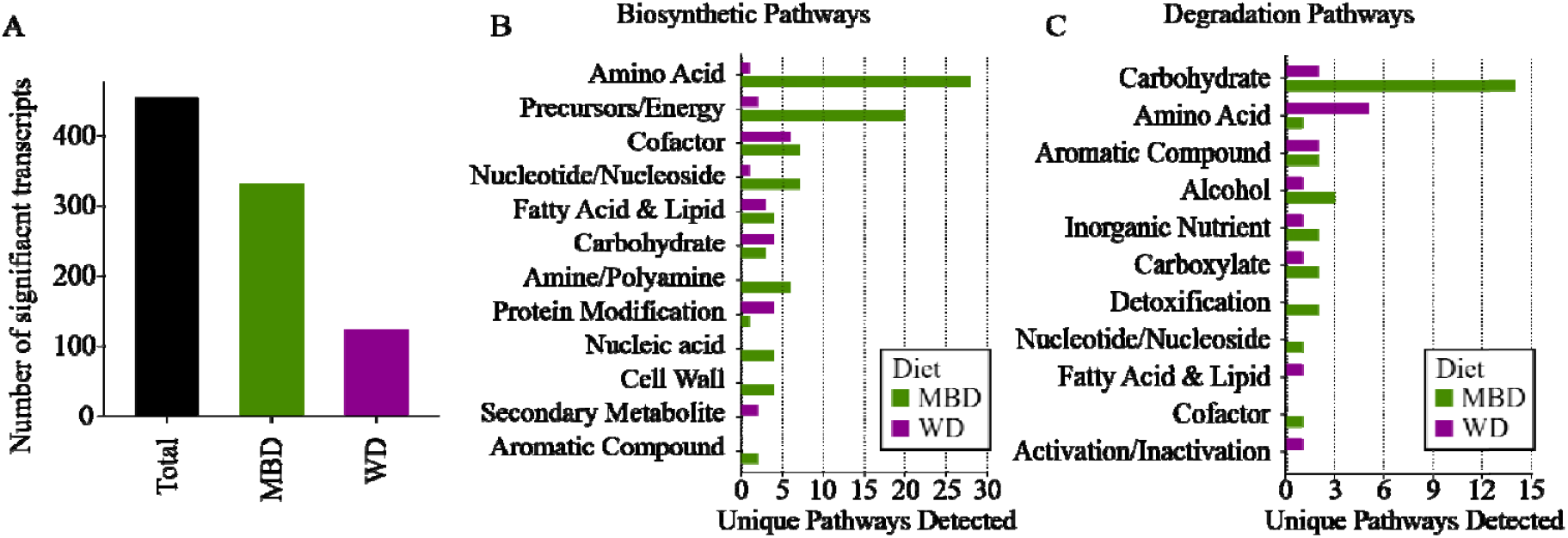
The number of significantly differentially abundant transcripts and the MetaCyc pathways that the transcripts mapped to. A) The total significant differentially abundant transcripts and the number of total significant differentially abundant transcripts in each diet. B) Number of unique reconstructed MetaCyc biosynthetic pathways found in each diet. C) Number of unique reconstructed MetaCyc degradation pathways found in each diet. All transcripts included had an unadjusted P value of ≤ 0.05, with a Benjamini-Hochberg adjusted P value ≤ 0.25.

Figure 2B shows that the gut microbiota on the MBD and WD were enriched in transcripts for functionally divergent biosynthetic pathways. The MBD was enriched in many biosynthetic pathways: e.g., biosynthesis of cofactors, amino-acids, nucleosides and nucleotides, cell wall components, fatty acids and lipids, and precursor molecules and energy. Conversely, on the WD the gut microbiota was enriched in pathways for the biosynthesis of carbohydrates, protein modifications, and secondary metabolites. Microbiota in the MBD contained more pathways for degradation of a broad variety of substrates than did the WD, as shown in Figure 2C. For example, on the MBD pathways to degrade carbohydrates, aromatic compounds, and alcohols were enriched, while on the WD only pathways for amino-acid degradation were enriched.

The metabolic differences in the pathways enriched in each diet (**Figures 2B** and **2C**) support that the MBD provided a fiber-rich nutrient source for the gut microbiota, which then utilized these sources for growth. This aligns with our prior fecal 16S rRNA gene copy number data, which we used as a proxy for biomass growth [27] that was significantly higher on the MBD compared to the WD. That pathways for amino-acid degradation were enriched in the microbiota of the WD suggests an environment in which polysaccharides were limited, available carbohydrates were quickly fermented, and amino acids became an important source for nutrition. Bacteria generally consume protein once carbohydrates have been exhausted[66,67], and studies have shown that increased intake of polysaccharides reduces protein fermentation even without changing protein intake [68].

### Microbiota expressed CAZyme transcripts for dietary carbohydrates in the MBD and host-glycans in the WD

One of the key differences between the two diets was carbohydrate composition. The total percentage of daily energy from carbohydrates remained the same in both diets [27,44], but the MBD had more energy from fiber and resistant starch (25.8 g of fiber/1000 kcals and 10.3 g of resistant starch/1000 kcals, respectively, compared to 6.5 g/1000 kcals and 1.2 g/1000 kcals for the WD). Because of this difference, we investigated the abundance of CAZyme transcripts, which are crucial for the metabolism of various carbohydrates, and found that their abundance (**Figure 3**) is influenced by dietary carbohydrate composition[33].

**Figure 3.**
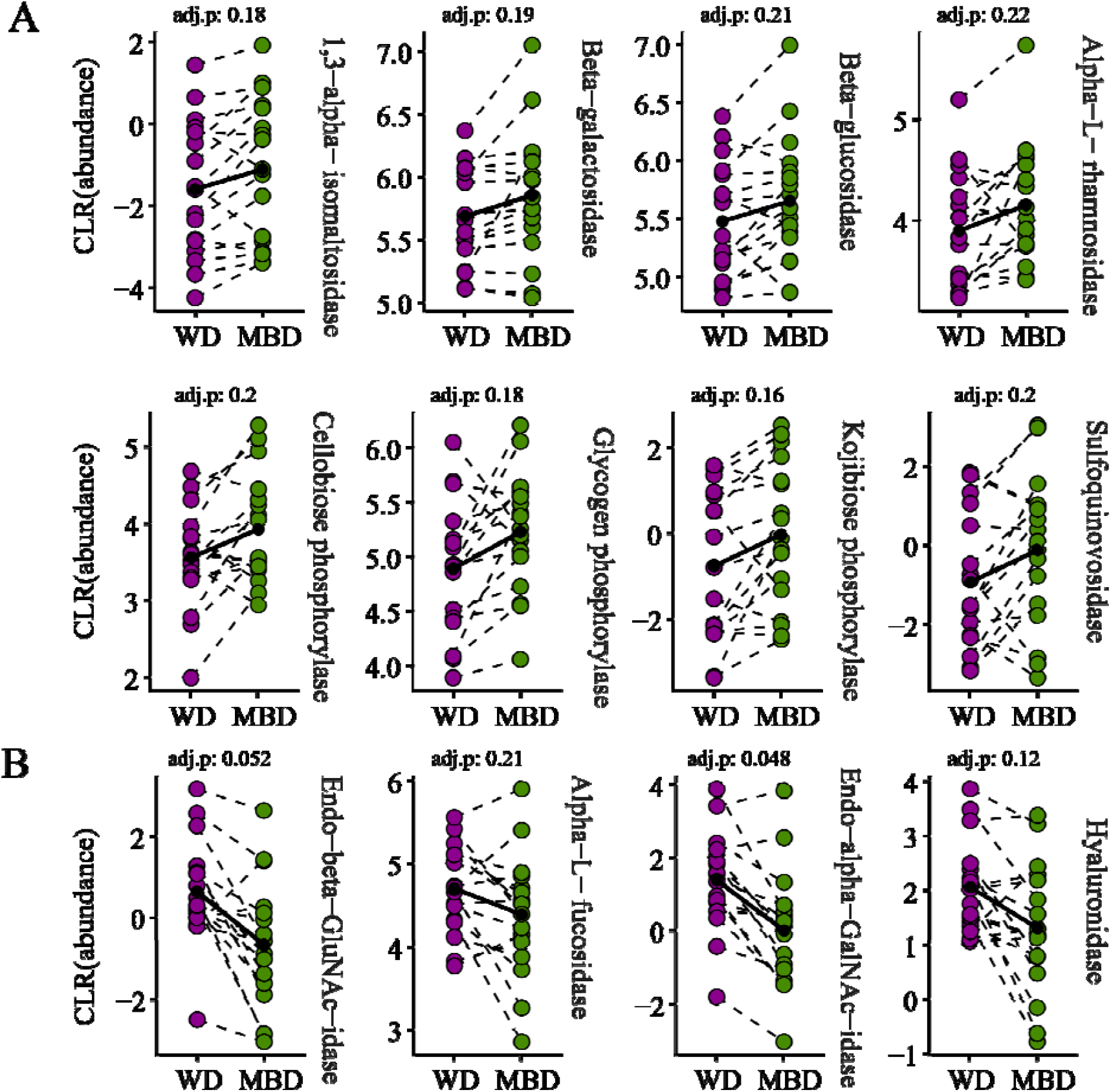
Differentially abudant CAZyme transcripts in the gut micorbiota during MBD and WD. A) CAZyme transcriptsthat were differentially more abundant on the MBD. B) CAZyme transcripts that were differentially more abundant on the WD. CLR: Centered log-ratio, GluNAc: N-acetylglucosamine, GalNAc: N-acetylgalactosamine. All significant transcripts had an unadjusted P value of ≤ 0.05, with a BH adjusted P value ≤ 0.25.

The microbiota in the MBD were enriched in transcripts for CAZymes that degrade dietary polysaccharides (**Figure 3A**). CAZymes like 1,3-α-isomaltosidase and glycogen phosphorylase degrade resistant starch [69–71]. Also abundant were transcripts for CAZymes that degrade hemicellulose like α-L-rhamnosidase, β-galactosidase, β-glucosidase, and cellobiose phosphorylase, the latter of which degrades cellobiose, a cellulose-derived disaccharide. The MBD gave an enrichment in sulfoquinovosidase, a CAZyme that hydrolyzes sulfoquinovoside, the sugar component in the plant lipid sulfoquinovosyl glycerol[72]. Finally, kojibiose phosphorylase, a CAZyme that degrades rare kojioligosaccharides, was also enriched on the MBD [73].

The WD microbiota were enriched in transcripts CAZymes that degrade host-derived carbohydrates, namely mucin and cell-bound glycoproteins (**Figure 3B**). Endo-α-N-acetylgalactosaminidase (endo-α-GalNAc-aminidase) cleaves the glycans from the polypeptide backbone of mucins, making the glycans available for other bacteria [74]. The α-L-fucosidase hydrolyzes the fucose sugars, one of the main components of mucins and other glycoproteins [75]. Similar to the endo-α-GalNAc-aminidase, the endo-α-N-acetylglucosaminidase (endo-α-GluNAc-aminidase) cleaves the glycans from host-cell membrane-bound glycoproteins [76]. The WD was also abundant in transcripts for the CAZyme hyaluronoglucosidase that hydrolyzes glycans commonly found in mammal cell-bound glycoproteins and the extracellular matrix found on the outside surface of mammal cells [77,78]. A metabolite analysis of the same fecal samples from this study by Igudesman *et al*. revealed increased fecal concentrations of key mucin components such as fucose, N-acetylglucosamine/N-acetylgalactosamine, and N-acetylneuraminate in the WD [79], indicative of enhanced mucin degradation [79].

The differences in abundances of CAZyme-transcripts between the microbiota for the MBD versus the WD highlight the effect of dietary macronutrient composition on the gut microbiota. The MBD delivered a plentiful source of polysaccharides to the microbiota to consume, while those polysaccharides were much lower in the WD, likely providing far fewer fermentable carbohydrates to the microbiota. We interpret that, for the WD, the microbiota had to turn to alternative, non-dietary carbohydrate sources, such as host glycans [25]. This pattern aligns with earlier findings from gnotobiotic mouse studies: A diet deficient in fiber drove the microbial community to scavenge host glycans in the colonic mucus layer [19], potentially compromising the gut barrier [23]. The increased fecal concentrations of mucin byproducts (e.g., fucose, N-acetylglucosamine/N-acetylgalactosamine, and N-acetyneuraminate) further support the notion that the WD environment favored host glycan turnover.

### Microbiota expressed protein-degrading transcripts for growth functions during the MBD, but nutrition and possible mucin-layer colonization with the WD

Although gut microbiota will generally first consume carbohydrates, protein is important as a source of amino acids, nitrogen, and energy. Protein degradation usually occurs in the distal colon, where the pH is near neutral. Although protein degradation is important for a healthy microbiota, it can produce compounds known to have detrimental effects on gut and human health. As diet composition is known to affect microbial protein degradation, we investigated differentially abundant transcripts for enzymes responsible for protein degradation, with the results summarized in **Figure 4**.

**Figure 4.**
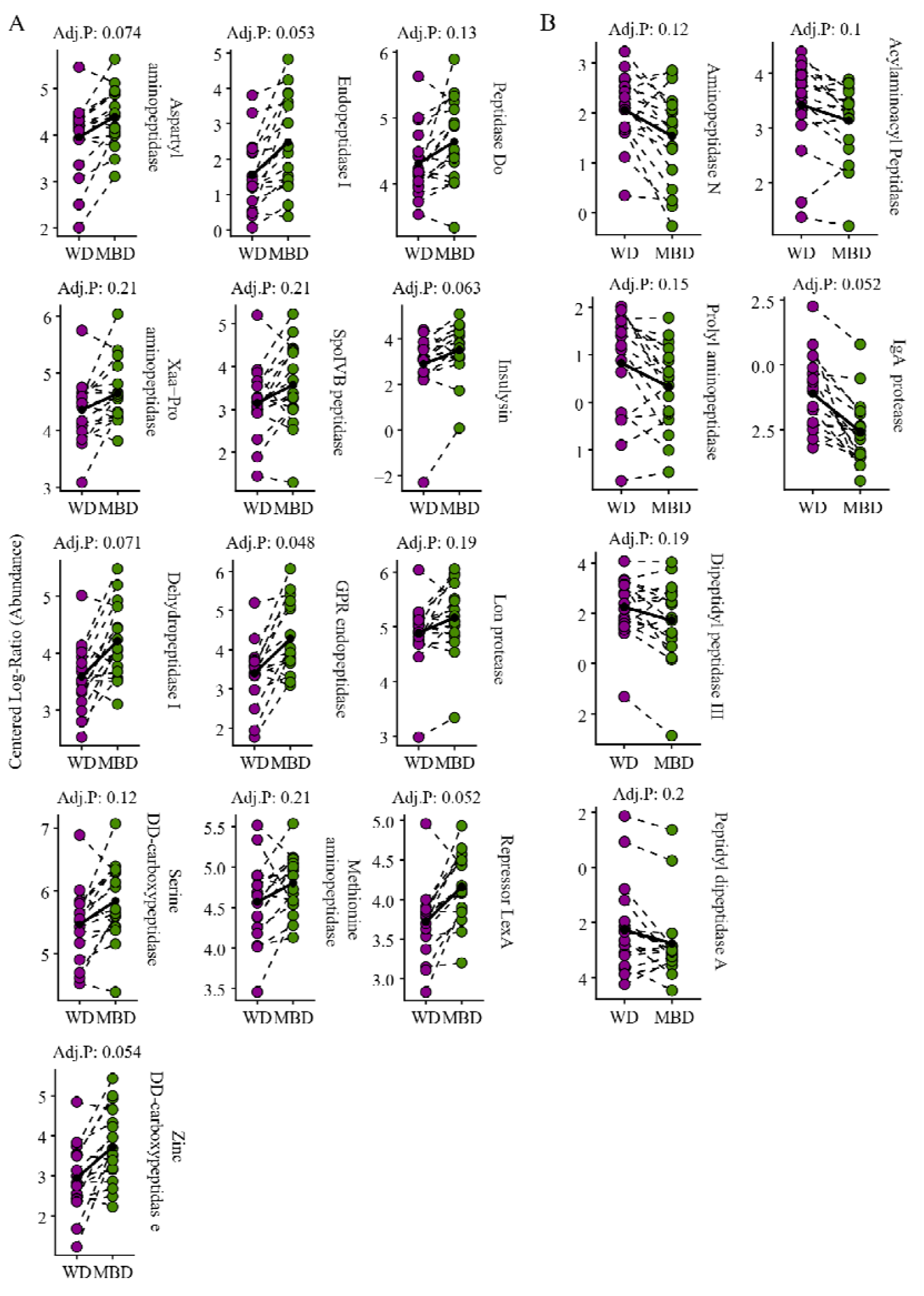
Differentially abudant protease transcripts in the gut micorbiota during WD. A) Proteases that were differentially more abundant on the MBD. B) Proteases that were differentially more abundant on the WD. CLR: Centered log-ratio. All significant transcripts had an unadjusted P value of ≤ 0.05, with a BH adjusted P value ≤ 0.25.

The microbiota with the MBD had more differentially abundant proteases than for the WD (**Figure 4A**), and the MBD’s proteases had a wider diversity of function than those under the WD. The MBD microbiota had more abundant transcripts for aspartyl- and Xaa-Pro aminopeptidases, along with dehydropeptidase I, all used for intracellular protein degradation. The MBD microbiota also was abundant in transcripts for serine and zinc dd-carboxypeptidases, used for cell wall formation, as well as endopeptidase I, spoIVB peptidase, and GPR endopeptidase, which are important for sporulation. Transcripts for methionine aminopeptidase, an enzyme that releases the initiator methionine from newly synthesized proteins, were higher under the MBD. Transcripts for degradation of denatured, misfolded, or aggregated proteins e.g., peptidase Do, lon protease, and insulysin, were more abundant under the MBD. Peptidases important for gene expression regulation, like lon protease and repressor LexA, were abundant for the MBD. For the WD (**Figure 4B)**, most of the abundant transcripts were for peptidases involved in protein turnover, including aminopeptidase N, prolyl aminopeptidase, dipeptidyl-peptidase III, and peptidyl-dipeptidase A. Transcripts for Acylaminoacyl-peptidase, an enzyme that degrades denatured proteins, were more abundant under the WD. With the WD, transcripts for IgA protease, which hydrolyzes IgA, an important antibody, were higher than with the MBD.

The diversity of peptidase functions for the MBD suggests that the microbiota were active in a range of catabolic and anabolic processes, such as building cell walls, synthesizing new proteins, disposing of misfolded proteins, and regulating gene expression. While those activities were still occurring under the WD, the WD’s microbiota seemed to be more focused on protein degradation. Analyses of fecal metabolites by Igudesman et al. revealed that amino-acid metabolites was enriched for the WD as compared to the MBD, such as the branch-chain fatty acids valine, leucine, and isoleucine, supporting this interpretation [79]. Additionally, the higher abundance of transcripts for a protease that hydrolyzes IgA, an antibody that modulates mucin-associated microbiota to maintain separation between microbiota and intestinal barrier [80], is important for many pathogens that colonize mucosal surfaces in the human body [81,82] and suggests that the microbiota under the WD were utilizing the mucin layer of the gut [81,82].

### Transcripts indicated that MBD microbiota fermented amino acids to produce biogenic amines, while WD microbiota fermented amino acids to produce uremic toxins

Both diets expressed transcripts important for protein degradation; however, this does not necessarily mean an increase in net proteolysis. Proteolysis is governed by a number of different factors, such as co-regulation [83], metabolic flux [84], and pH [85]. Given this, we looked at differentially abundant transcripts involved in amino-acid fermentation. As shown in **Figure 5**, transcripts for enzymes that ferment amino acids were differentially enriched in each diet in ways that have potential implications for host health.

**Figure 5.**
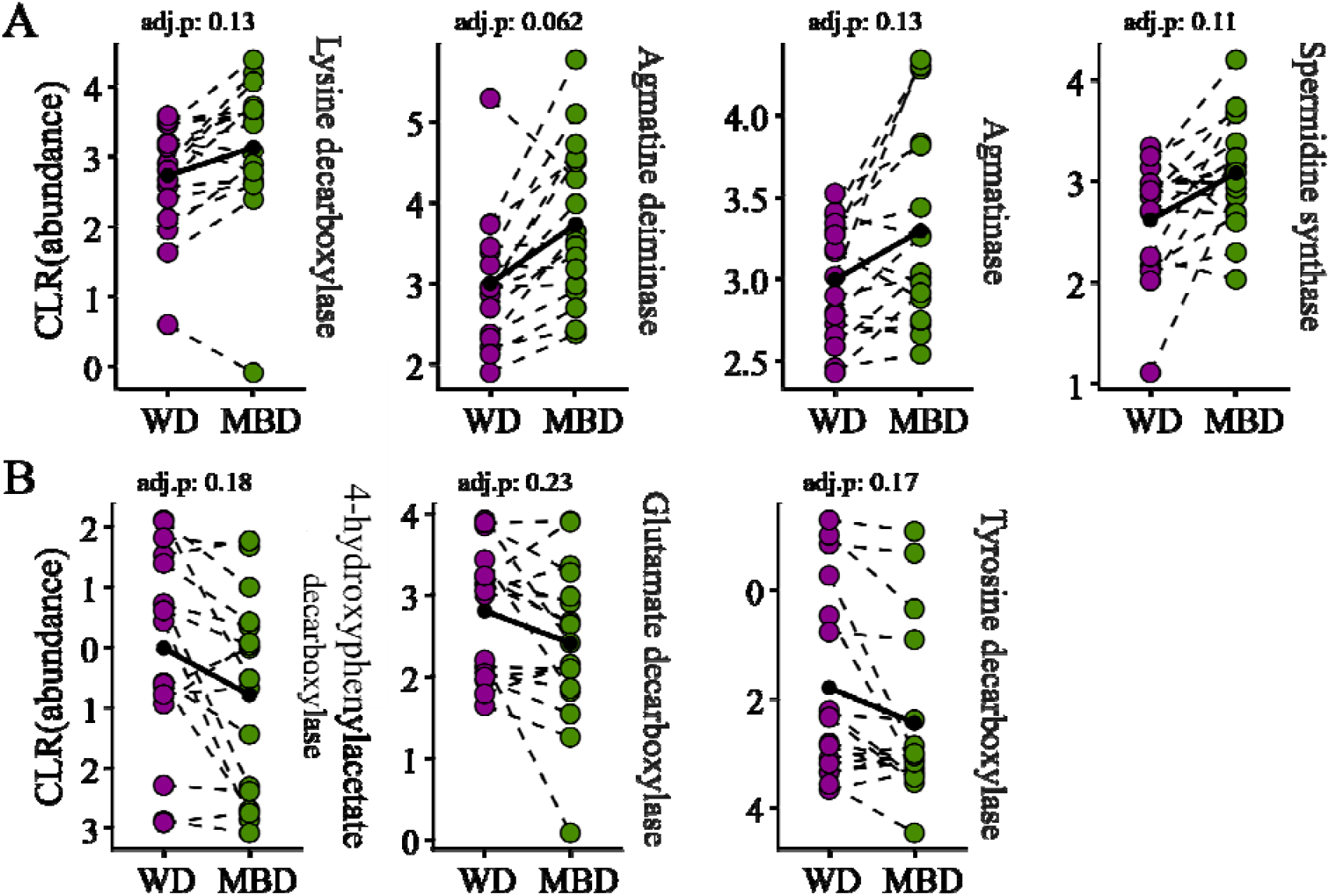
Differentially abundant microbial transcripts encoding for amino acid-degrading enzymes. A) Differentially abundant transcripts for amino acid degrading enzymes in MBD. B) Differentially abundant transcripts for amino acid degrading enzymes in WD. CLR: Centered log-ratio. All transcripts had an unadjusted P value of ≤ 0.05.

The WD microbiota were enriched in transcripts for 4-hydroxyphenylacetate decarboxylase, glutamate decarboxylase, and tyrosine decarboxylase; 4-hydroxyphenylacetate decarboxylase produce p-cresol from tyrosine [86], glutamate decarboxylase produce γ-aminobutyric acid (GABA) from glutamate [87], and tyrosine decarboxylase produces tyramine from tyrosine [88] (**Figure 5A)**. In contrast, the MBD microbiota were enriched in transcripts for lysine decarboxylase, agmatinase, agmatine deiminase, and spermidine synthase; lysine decarboxylase produces cadaverine from lysine, agmatinase and agmatine deiminase produce putrescine from agmatine, and spermidine synthase produces spermidine from putrescine (**Figure 5B**).

Fermentation of amino acids released during protein degradation can produce metabolites that have implications for host health: e.g., uremic toxins [89], neurotransmitters [90], and biogenic amines [91]. The 4-hydroxyphenylacetate decarboxylase and tyrosine decarboxylase in the WD microbiota ferment aromatic amino acids to produce the uremic toxin p-cresol and the biogenic monoamine, tyramine. Both metabolites have been linked to gastrointestinal disorders. Fecal tyramine has been found to be elevated in people suffering from IBS [92] and p-cresol has been linked to Crohn’s diseases and IBD. [93] Tyramine has been shown to disrupt tight junctions *in vivo* in zebrafish [94] and impair DNA repair and fatty acid β-oxidation in HT29 cell cultures [92]. Like tyramine, p-cresol also damages the junctions between epithelial cells and weakens the intestinal barrier [95]. P-cresol has been shown to interfere with neurotransmitter metabolism and impair mitochondrial function [96]. GABA, the neurotransmitter produced by glutamate decarboxylase, is responsible for modulating mood, anxiety, and stress response [97].

The three transcripts enriched in the microbiota for the MBD produce polyamines: cadaverine, putrescine, and spermidine, which are important for maintaining a functional gut barrier [98,99]. Analysis of fecal metabolites by Igudesman et al. showed elevated spermidine, p-cresol, and 4-hydroxyphenylacetate, the p-cresol precursor for the WD, but none under the MBD [79]. The lack of these fermentation products with the MBD was likely due to the increased amount of carbohydrates coming from fiber and resistant starch in the MBD. Increased fiber and resistant starch provide microbes with carbohydrates and reduce protein fermentation in the distal colon [100]. For example, resistant starch reduced p-cresol detected in mice fed a diet supplemented with tyrosine [101]. Additionally, microbes themselves may even modulate host polyamine metabolism. A study showed that a lactobacillus strain introduced into colitis-induced mice increased expression of polyamine degrading enzymes in the host which resulted in reduced polyamine concentration in the gut [102].

## Conclusion

In summary, this study demonstrates that, while the overall functional diversity of the gut microbiota remained relatively stable at the transcript level, diets that differed in particle size and resistant starch and fiber content shifted the types of metabolic processes being actively expressed. The relationships are illustrated in **Figure 6**. The MBD promoted a resource-replete state that supported robust biosynthetic and carbohydrate-fermenting pathways, while the WD led to a resource-limited state marked by elevated host-glycan and protein degradation. The MBD supported an ecosystem geared toward carbohydrate fermentation and biosynthesis, presumably benefiting host health via enhanced SCFA production and reduced reliance on host glycan degradation. In contrast, the WD’s reduced fiber input fostered increased mucin and protein breakdown pathways, yielding metabolites that may adversely affect the gut barrier and systemic metabolism.

**Figure 6.**
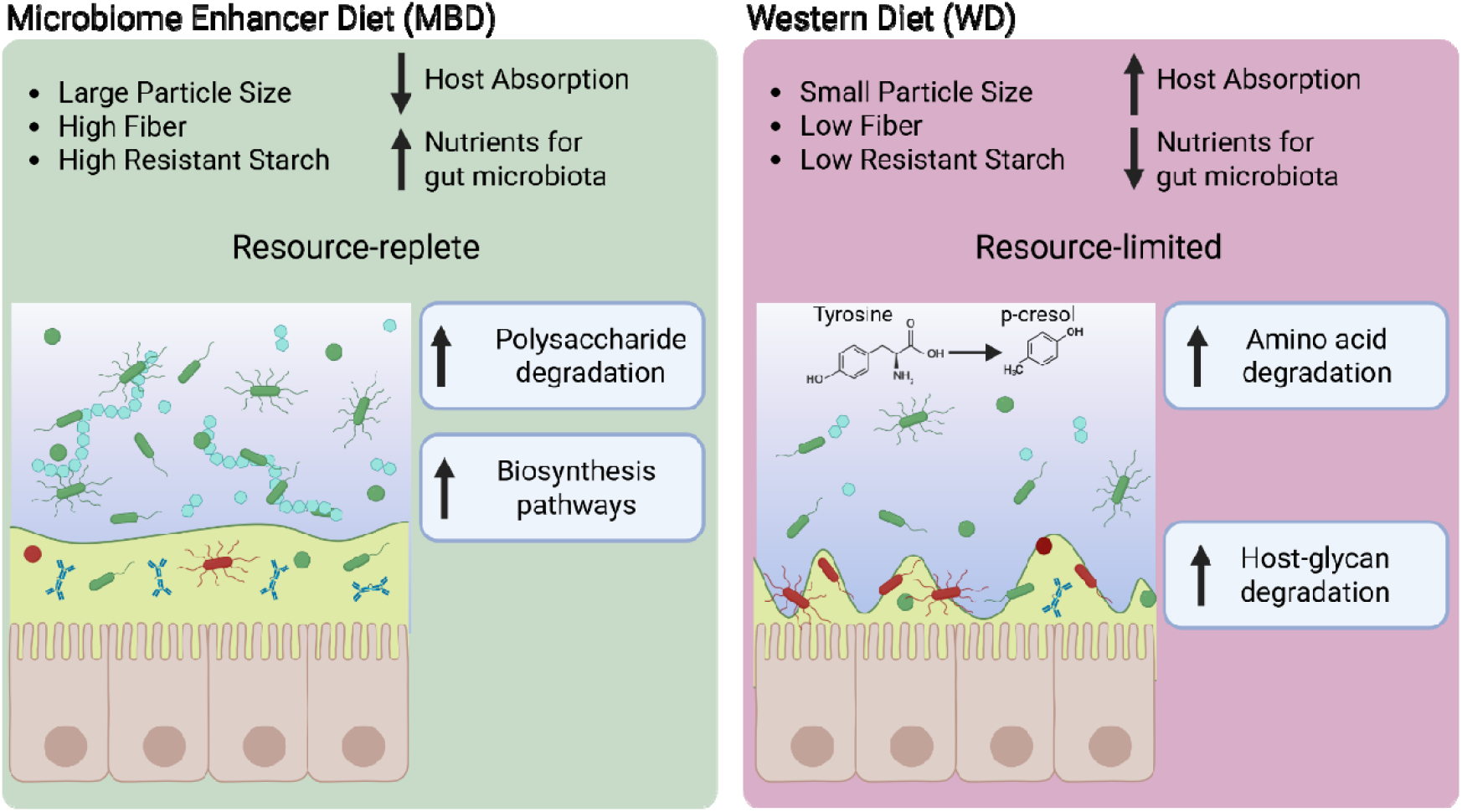
Conceptual overview of the microbial functional response to the Microbiome Enhancer Dier (MBD) and Western Diet (WD). Each diet contained the same proportion of carbohydrates but differed in the characteristics of those carbohydrates. In the MBD, the carbohydrates had larger particle size and higher fiber and resistant starch content. In the WD, the carbohydrates had smaller particle size and lower fiber and resistant starch content. Macronutrients in the MBD were less absorbable by the host than in the WD, which delivered more fermentable carbohydrates to the gut microbiota than the WD. This made the gut microbiota “resource-replete” with the MBD and “resource-limited” with the WD. The gut microbiota responded to this difference in resource availability with increased transcription for carbohydrate active enzymes (CAZymes) degrading polysaccharide and increased transcription for proteins contributing to biosynthesis pathways in the MBD. In contrast, the gut microbiota with the WD responded to limited resource availability by increasing transcription for amino acid degradation and CAZymes degrading host-glycans such as mucin. *Figure created using Biorender*.

Some limitations of our study should be noted. First, our limited sample size necessarily means that these results are exploratory in nature, which allow us to generate hypotheses that need to be validated in future studies. Although our controlled feeding design enhanced precision in dietary interventions, the relatively short intervention period may not fully capture the long-term effects of sustained dietary changes. Indeed, previous longer-term observational studies (e.g., ∼6 months) by Mehta and colleagues noted significant intra-individual shifts in the fecal metatranscriptome [103]. Larger sample sizes or longer-term studies could yield additional insights, especially regarding how enduring these functional shifts might be overtime. Second, metatranscriptomics offers a valuable snapshot of gene expression, but does not capture post-translational modifications or protein activity directly. Integrating metaproteomic data in future studies could further illuminate the functional repercussions of these altered pathways. Lastly, although we controlled for total macronutrients, the inherent complexity of whole foods means that specific phytochemicals, fiber types, and other bioactive compounds could also influence the microbiota’s functional responses.

These findings highlight the power of metatranscriptomics for revealing intriguing hypotheses about the mechanistic underpinnings of how dietary patterns influence the gut microbiota’s functional repertoire, which may ultimately shape health trajectories. Future work that confirms and extends these observations to broader populations, diverse dietary regimens, and longer intervention periods can deepen our understanding of how best to modulate gut microbiota through precision nutrition strategies.

## Author Contributions

BD and RK-B designed the study analysis. BD performed bioinformatic and statistical analyses. SRS, RK-B, and BER designed the parent clinical study and obtained funding. KDC and EAC supervised the clinical trial. AEM helped with writing. AEM and CMW provided critical feedback on the data. BD wrote the first draft of the manuscript with edits and input from all authors. All authors critically reviewed the manuscript, provided feedback, and agreed with the content.

## Nucleic Acid Sequences

DNA and RNA sequencing data from this study can be found in the BioProject database under the accession codes PRJNA913183 and PRJNA947193.

