## Supplementary Figure 1 for "Microbial Feast or Famine: dietary carbohydrate composition and gut microbiota metabolic function"

**Supplementary Material**

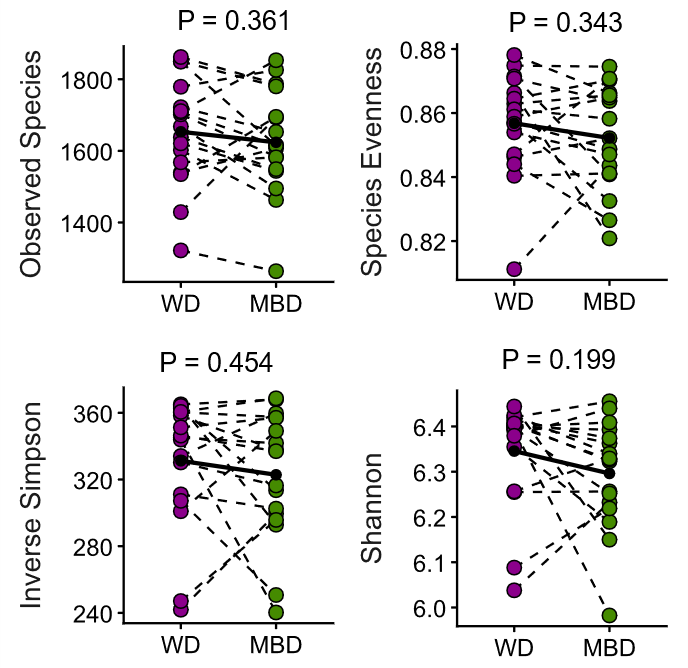

**Supplementary Figure 1. Alpha-diversity metrics.** Measured alpha-diversity metrics observed species, evenness, Inverse Simpson, and Shannon diversity were not significantly different between the WD and the MBD. N = 17.
